# Alarm tones, music and their elements: A mixed methods analysis of reported waking sounds for the prevention of sleep inertia

**DOI:** 10.1101/607358

**Authors:** Stuart J. McFarlane, Jair E. Garcia, Darrin S. Verhagen, Adrian G. Dyer

## Abstract

Sleep inertia is a potentially dangerous reduction in human alertness and occurs 0 – 4 hours after waking. The type of sound people set as their alarm for waking has been shown to reduce the effects of sleep inertia, however, the elemental musical factors that underpin these waking sounds and their relationship remain unclear. The goal of this research is to understand how a particular sound or music chosen to assist waking may counteract sleep inertia, and more specifically, what elements of these sounds may contribute to its reduction using a mix methods analysis. Through an anonymous, self-report online questionnaire, fifty participants (*N = 50*) reported attributes of their preferred waking sound, their feeling towards the waking sound, and symptoms of sleep inertia after waking. This data enabled the analysis and comparison between these responses to define statistically significant interactions. Our results show that there is no significant relationship between sleep inertia and the reported waking sound, nor the subject’s feeling towards this sound. However, we found that the melodicity of a chosen waking sound does effect sleep inertia. A sound that is perceived as melodic, produces less sleep inertia in comparison to a sound considered to be neutral (neither unmelodic nor melodic). Furthermore, a secondary analysis reveals that this is an important factor for waking stimulus design as it suggests that the amount of perceived rhythm will affect the perception of melody, and in turn, may influence the severity of sleep inertia on a secondary level. Our results reveal that the inclusion of detailed descriptive terms (musical elements) in addition to macro classifications (e.g. “pop music”) for stimulus testing would benefit future research and our understanding of waking audio’s effects on sleep inertia.

## Introduction

> *“The morning started disastrously. I slept through two alarms, one set for 0600 and another a half-hour later to remind me to take some CEO pictures. My body apparently went on strike for better working conditions.”*
>
> **NASA astronaut journal report during orbit aboard the International Space Station. [1]**

Sleep inertia (*SI*) is a sleep-wake disorder characterized by low arousal and reduced cognition [2]. Initiated upon waking, *SI’s* symptoms can last for seconds, minutes or hours, where extended *SI* may impact human performance in a variety of fields and occupations [3–9]. This has been highlighted as a likely factor in the 2010 Air India Express air crash disaster that resulted in 158 fatalities. It has been shown that the captain of the aircraft had recently woken from an in-flight nap just prior to the crash. The poor decisions made after napping were attributed to the disaster, and have been linked to the effects of *SI* [5].

There has been growing research interest into the mechanics and architecture of *SI* [10–15], however the modalities and means to activate waking remains in its early stages with respect to this issue. YouGov [16] report that out of 586 participants surveyed, 68.2% use a form of alarm for waking, of these, 23% use an alarm clock, 14.9% a clock radio, and 26.3% an alarm on a cell phone. These figures show alarms are still an important means to wake up, and given the 24-hour society in which we often live and work, the need for peak performance from our waking device, and the stimuli they produce, is advantageous to counteract the negative effects of *SI*.

Research in the field of human factors and psychology has provided initial insights into the application of countermeasures to reduce *SI*. Countermeasures are strategies to be implemented upon waking as opposed to methods that may consider circadian management, or pre-sleep hygiene techniques (routines to assist and promote sleep). Within the current literature investigating experimental *SI* countermeasures include light, temperature and sound [11, 17–19]. In the context of our research, two previous studies present findings on the effects of sound as a countermeasure for *SI*. Tassi et al [11] concluded that noise can reduce *SI* when deployed as an intense waking alarm [11], while Hayashi et al [19] discovered that ‘high-preference popular music’ as chosen by participants has the potential to reduce the intensity of *SI* after a short nap. Both studies support the use of sound and music as a countermeasure for *SI*, although many questions remain as to the auditory mechanics and aesthetics required for best practice design of such stimuli, hence, a consistent approach to minimizing *SI* through audio is yet to be determined.

Musical elements are components of music or sound that exist to assist in the description and production of music. These elements can be treated individually or combined to produce an infinite array of musical aesthetics, effects and compositions. Musical elements include, yet are not limited to, melody, rhythm, pitch, tempo and volume. These elements differ from sound types and genres as they afford the description of audio through musical characteristics and have the capacity to expose factors which may be neglected in sound type classification [20, 21].

For one example, in the context of this research field, a subject may describe auditory stimuli as ‘popular music’, however, when reported as musical elements, the audio can be defined as ‘very melodic’, ‘rhythmic’, ‘high pitch’, ‘fast tempo’, and ‘high volume’. From these descriptions, ‘pop music’ may now be analyzed and understood through musical terms to establish in greater detail what the respondent may actually be hearing, regardless of the genre or stimulus title.

This mixed method research explores waking audio and its effects on *SI* through the deployment of a self-reporting online questionnaire designed to answer three primary research questions, (i) ‘*Do waking sound types counteract the effects of SI?’*, (ii) ‘*Do subjective feelings towards waking sound types counteract the effects of SI?’*, and (iii) ‘*Do the musical elements of waking sound counteract the effects of SI?’*.

To achieve our objectives, several separate analyses where performed comparing the respondents’ reported intensity of *SI* against their waking sound types, subjective feelings, and the musical elements of their waking audio. Additionally, further analysis investigates secondary level attributes of waking types and musical elements with respect to *SI*. Lastly, response time evaluation and alertness factors are evaluated, together with qualitative observations.

## Materials and Methods

### Ethics statement

All research methods and data collection were approved by RMIT’s University College Human Ethics Advisory Network (CHEAN) (ref: CHEAN A 201710-02-17). The participants consisted of post-graduate students and staff members from the schools of Media and Communications, Industrial Design, and the School of Health and Biomedical Sciences, as well as the Australian Sleep Association. All respondents consented to participate by completing the online study. This was stipulated to the subjects in the ‘Invitation to participate’ email distributed during the recruitment period. Potential participants were encouraged to undertake the study without bias towards music aptitude or waking method. All respondents (See Results) were 18 years of age and above consisting of males and females. Consistent with ethics, age and gender are for demographics only and are not subject to analysis in this study.

### Data collection

Data was collected via a self-report online questionnaire. The submitted data was captured digitally via the use of the online software system Qualtrics [22], where the questionnaire was contained and operated. Qualtrics is software specifically created for the undertaking of online questionnaires and surveys enabling researchers to design and implement their studies for ethically compliant distribution and data collection. The data obtained by Qualtrics is securely stored and only available for download and analysis by researchers with appropriate clearance.

### Questionnaire – Development and Description

We designed a custom questionnaire which incorporated 17 items and 4 open ended response prompts. The questionnaire used a combination of Likert scale, multiple choice and open-ended questions, to return qualitative and quantitative responses. The design comprises of four sections: Welcome Page (**Section 1**), Demographic Information (**Section 2**), Waking Sound (**Section 3**), and Sleep Inertia (**Section 4**), in total, requiring approximately ten minutes to complete. The ten-minute data collection was designed based on pilot results to minimize disruption to each subject during their natural ‘day-to-day’ waking routine, so as to maximise the ecological validity of the result.

Section 1 is an introduction to the study and clarifies the participants’ obligations prior to completing the task. This includes a reminder that by completing and submitting the survey the participant has given their specific consent to partake in the study (S 1. Fig 1).

Section 2 (Items 1-5, S 2. Fig 2) is comprised of Likert scale (unipolar Item 5, bipolar Item 4) and multiple-choice (Items 1-3) questions, which gather the demographic data of the respondents, their music appreciation, and musical aptitude. This data is paired with responses from Section 3 (Items 7-9, 12, Fig 1), and reported in the results under the following categories: Respondents (Table 7), Music Appreciation & Aptitude (Table 8), and Alarm Adoption & Application (Table 9).

**Fig 1.**
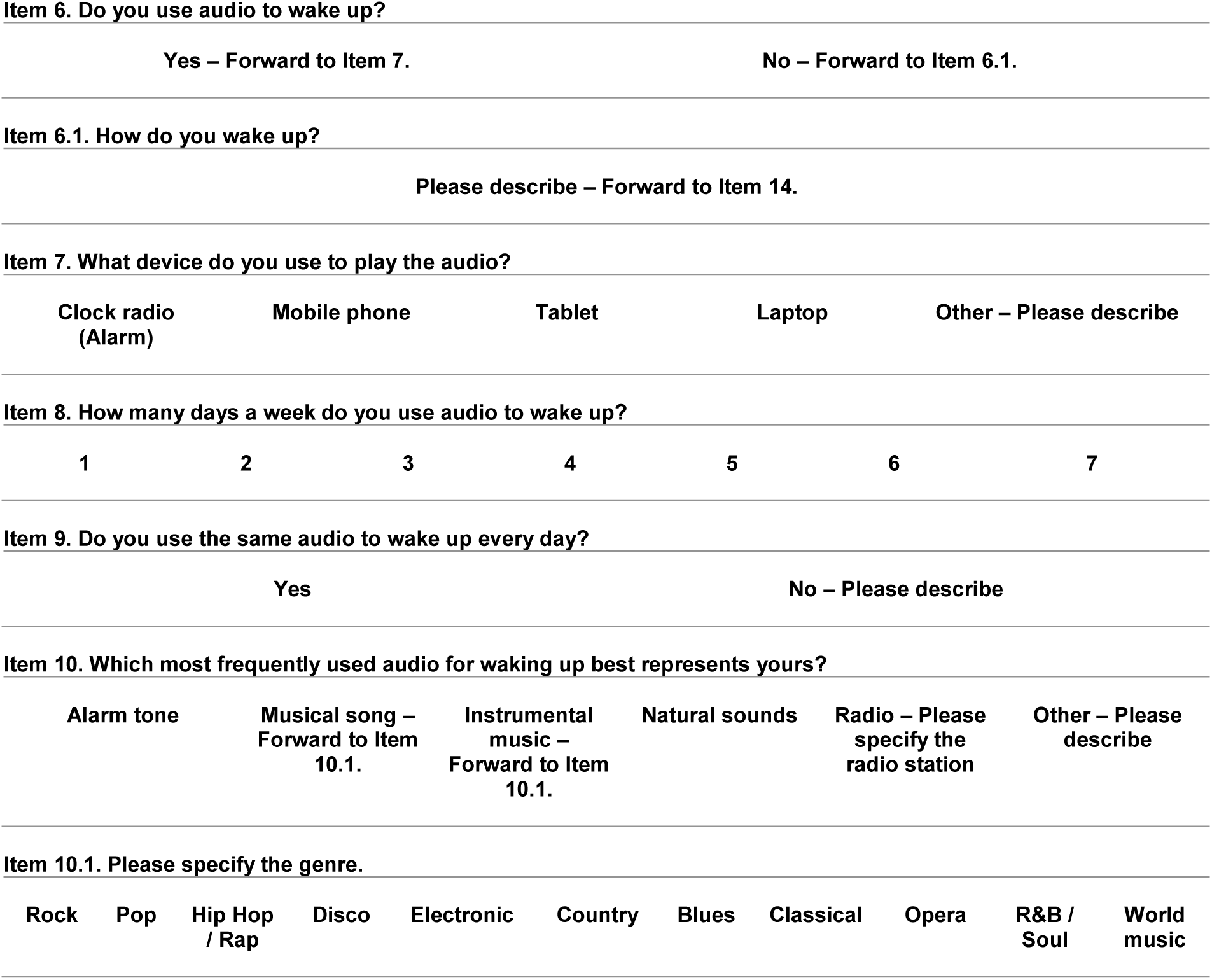
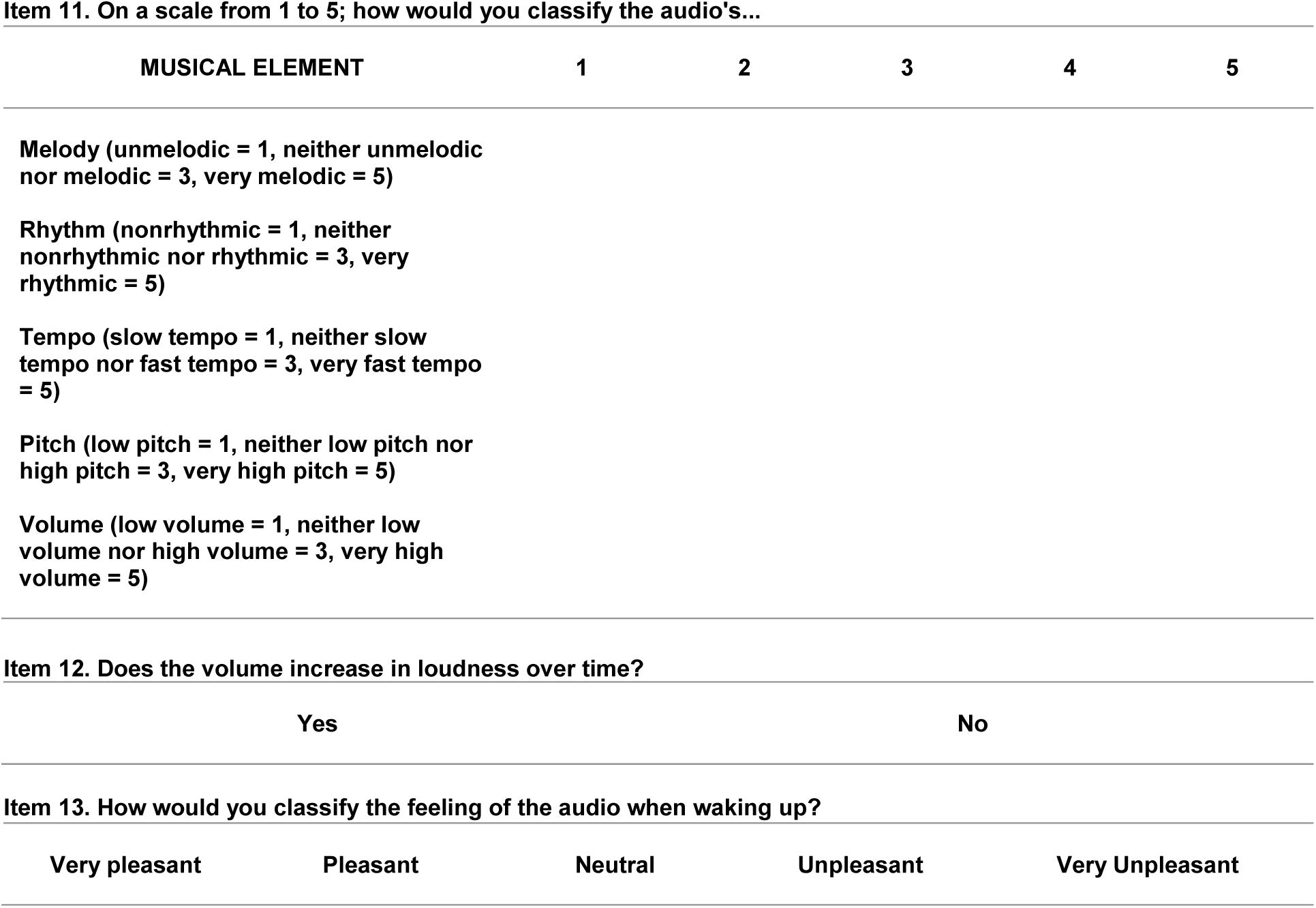
Waking sound & sleep inertia questionnaire Section 3.

Section 3 gathers information relating to the respondents use of audio for waking (if applicable), and the reported musical detail of the audio. This section incorporates Likert scale, multiple choice and open-ended questions. See Figure 1.

The first item in Section 3 (Item 6, multiple choice) identifies whether respondents use audio to assist them in waking. If the participant does not, they are prompted to describe their waking method, then forwarded to Item 14 in Section 4. If the participant does use audio, they are required to report what device is preferred to play the audio (Item 7, multiple choice), the number of days per week at which audio is employed (Item 8, multiple choice), and consistency of use (Item 8, multiple choice). When compiled, these items contribute to the participant profile (Alarm Adoption & Application, Table 9) previously outlined in Section 1’s description.

Item 10, ‘Which most frequently used audio for waking up best represents yours?’ requires the participants to nominate or specify the type of audio they use from six options provided. This item is the essential component to determine each participants’ waking sound type, and is elemental in formulating the analysis in response to primary research question (i) *‘Do waking sound types counteract the effects of SI?’*.

In designing the options for item 10, we first defined ‘sound type’ [23, 24] as the most suitable overarching terminology in this context to define all audio that may be reported as waking sound stimuli (e.g. musical genre, auditory tone, white or pink noise, human speaking, the sound of the wind, or aircraft engine noise). Secondly, the first five options for selection (Alarm tone, Musical song, Instrumental music, Natural sounds) were elected as they are familiar descriptions of audio sound types which respondents may use for waking. To determine the categories, we surveyed available pre-set sound types and custom audio functionality provided by several device manufacturers [25–31]. The sixth option (Other) allows for the respondent to describe their specific sound type if desired. In sum, these options allow for the breadth of potential waking sound types respondents may report.

Operationally, if ‘Musical song’ or ‘Instrumental music’ are selected, the respondent is then forwarded to Item 10.1 and are requested to specify which genre represents their waking sound type. The categories of genre for selection have been adapted from the Short Test of Music Preference (STOMP) [32]. Similarly, when ‘Radio’ is specified the respondent is prompted for the specific station. If ‘Other’ is selected, the participant is requested for a description.

Item 11 is another key factor of the study and is employed to analyse primary research question (iii) ‘*Do the musical elements of waking audio counteract the effects of SI?’*.

This item gathers each respondent’s classification of their waking audio’s musical elements which is used to establish a profile of the stimuli from a fundamental musical level. Each participant is required to rank the musical elements of their waking audio on a 5-point bipolar Likert scale that we developed. These specific ranks have been selected to afford descriptions of the participant’s waking audio musical elements through subjective interpretations (e.g. negative, neutral, positive). Ranks 1, 3, and 5 are labelled (e.g. 1 = unmelodic, 3 = neither unmelodic nor melodic, 5 = very melodic), while rank 2 and 4 remain uncategorized to reduce respondent bias. When reporting and discussing the results, we included labels to ranks 2 and 4 for continuity which we have defined in the rank response codes shown below (Table 1).

**Table 1.** Section 3 (Item 11) Likert scale rank response codes.

| REPORTED LIKERT SCALE RANK CATAGORIZATION & INTERPRETATION |  |  |  |  |  |
| --- | --- | --- | --- | --- | --- |
| MUSICAL ELEMENT | 1 | 2 | 3 | 4 | 5 |
| Melody | unmelodic | somewhat unmelodic | neither unmelodic nor melodic | melodic | very melodic |
| Rhythm | nonrhythmic | somewhat nonrhythmic | neither nonrhythmic nor rhythmic | rhythmic | very rhythmic |
| Tempo | slow tempo | somewhat slow in tempo | neither slow tempo nor fast tempo | fast tempo | very fast tempo |
| Pitch | low pitch | somewhat low in pitch | neither low pitch nor high pitch | high pitch | very high pitch |
| Volume | low volume | somewhat low in volume | neither low volume nor high volume | high volume | very high volume |

Section 3 is completed with Items 12 and 13. Item 12 (multiple choice) requests if the respondent’s waking audio rises in volume over time (a detail relating to contemporary sound design and increasingly common as an option available in many of today’s alarm devices and applications [25–29]). This item’s results are contained in the ‘Alarm adoption & application’ demography.

Item 13 investigates the respondents’ subjective feelings towards their waking sound and allows for the statistical analysis of primary research question (ii) ‘*Do subjective feelings towards waking sound types counteract the effects of SI?’*.

Deployed as a 5-point bipolar Likert scale, this design rates the participant’s subjective feeling towards their waking sound as: Very pleasant, Pleasant, Neutral, Unpleasant, and Very Unpleasant.

Section 4 is the leading component to the ‘Waking Sound and Sleep Inertia’ questionnaire as it enables the statistical reporting of respondents’ *SI* intensity after waking. Adapted from the Sleep Inertia Questionnaire (SIQ) developed by Kanady and Harvey [33], the SIQ has been researched and analysed to be a reliable measure of *SI* [33]. Our adapted SIQ begins with the question; ‘After you wake up, to what extent do you…’ [33] and is followed by each item. All respondents are required to rate each item as either, Not at all = 1, A little = 2, Somewhat = 3, Often = 4, All the time = 5. In the context of this study, eight items were removed from the original SIQ questionnaire as they are already included in other items in the questionnaire, or are not a focus of this study. See Table 2.

**Table 2.** Omitted items from the original SIQ.

| OMITTED ITEMS |
| --- |
| Need an alarm to wake up? |
| Notice that you get winded more easily? |
| Notice that you feel tense? |
| Feel anxious about the upcoming day? |
| Dread starting your day? |
| Wish you could sleep more? |

Section 4 is completed with three final multiple-choice items (15, 16, 17). When statistically analysed, these items test *SI’s* intensity with respect to time duration of perceived alertness, questionnaire completion, and time of day to complete the questionnaire. The results allow us to establish if the respondents *SI* intensity alters with respect to time. (Fig 2)

Item 14. After you wake up, to what extent do you…

**Fig 2.**
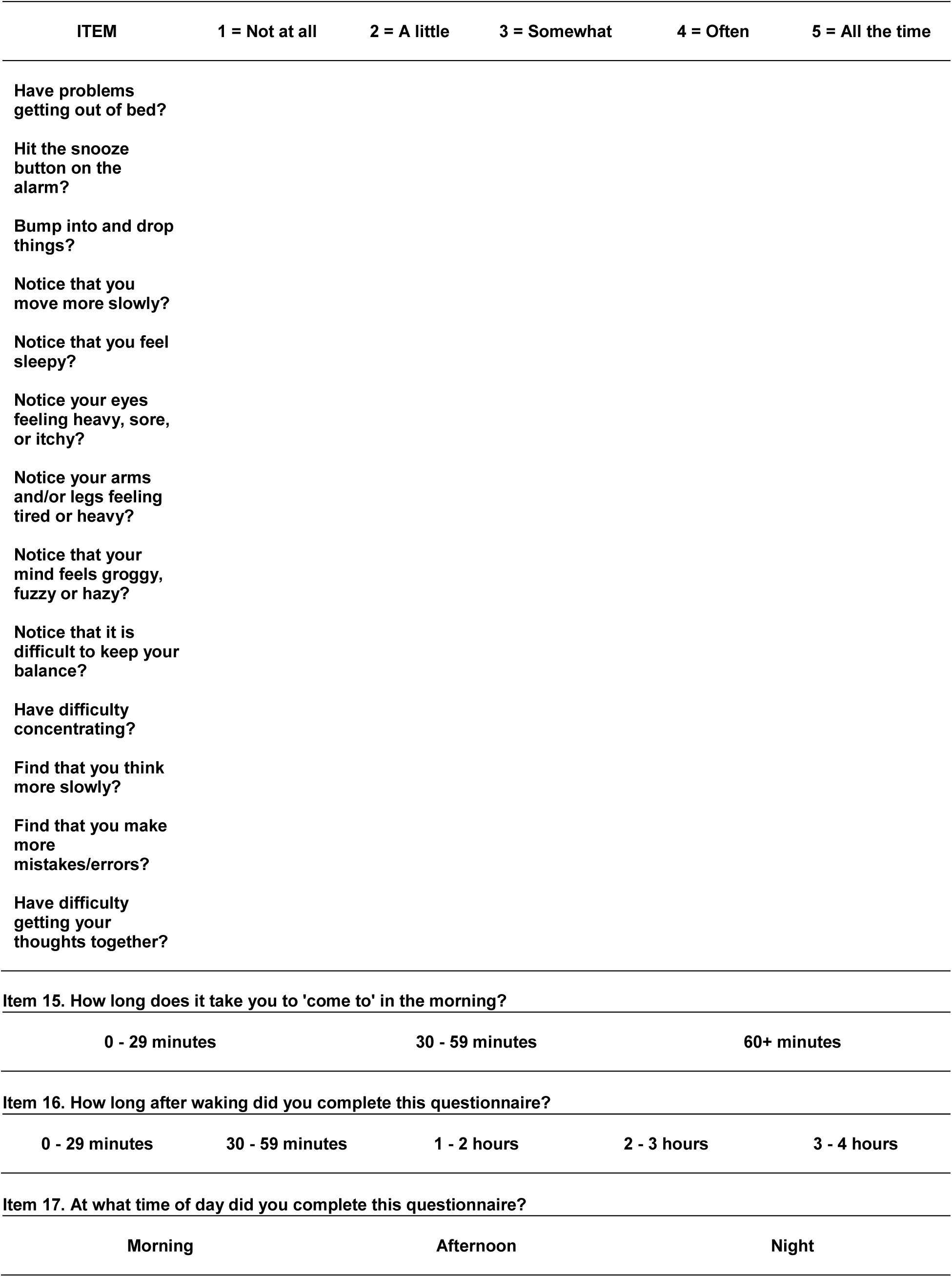
Waking sound & sleep inertia questionnaire Section 4.

### Procedure

The project was enabled through an anonymous online questionnaire that is performed upon waking by each participant. This method was chosen to maximise the natural contextual environment in which subjects use auditory alarms. Subjects were invited to participate via email through their respective schools or member association. This method for distribution was defined to ensure the anonymity of the participants. The contents of the email included an introduction, title and overview of the research, who is conducting the study, participants’ rights and responsibilities, instructions for how to undertake the questionnaire, and contact information for any further enquiries regarding the test. It is stipulated that the subjects are required to complete the questionnaire within four hours after waking, and that all questions must be responded to when prompted. Furthermore, the invitation email explicitly states that by completing the questionnaire, the participants are agreeing to undertake the research. By so doing, the invitation email effectively replaces a traditional participant information ‘hard copy’ form and serves as the online equivalent. The invitation to participate included a link to the study which directed each respondent to the online questionnaire for commencement. The study was launched during May 2017 and concluded in May 2018. A transcript of the invitation to participate email can be found in the supporting material section.

### Data processing

The total number of initial respondents was 83 (*N* = 83) which was filtered omitting any inconsistent or incomplete responses, reducing *N* to (*N* = 73). Further, this study requires the analysis of waking sound stimuli, therefore the ‘No waking sound’ responses were disregarded, reducing *N* to (*N* = 50). The *SI* intensity is determined as the mode of each participant’s response within the adapted SIQ questionnaire items in Section 4 (Fig 2). The raw data was initially filtered through Microsoft Excel [34], then imported to SPSS [35] for statistical analysis.

## Statistical analysis

### Primary analysis

To address the three primary research questions, (i) ‘*Do waking sound types counteract the effects of SI?’*, (ii) ‘*Do subjective feelings towards waking sound types counteract the effects of SI?’*, and (iii) ‘*Do the musical elements of waking audio counteract the effects of SI?’*, we conducted a series of test analyses. The results from the adapted SIQ (Item 14) are analysed against the data sets obtained from Items 10, 13, and 11 as described in Material and Methods. Table 3 contains the sequence of analyses, and each test we performed for the primary research questions.

**Table 3.** Primary test analysis.

| TEST ANALYSIS | PRIMARY RESEARCH QUESTION | ANALYSIS COMPARISONS |
| --- | --- | --- |
| No 1. | (i) *‘Do waking sound types counteract the effects of SI?’* | SI (Item 14) vs Sound type (Item 10) |
| No 2. | (ii) *‘Do subjective feelings towards waking sound type counteract the effects of SI?’* | SI (Item 14) vs Subjective feeling (Item 13) |
| No 3. | (iii) *‘Do the musical elements of waking audio counteract the effects of SI?’* | SI (Item 14) vs Music element - Melody (Item 11) |
| No 4. | (iii) *‘Do the musical elements of waking audio counteract the effects of SI?’* | SI (Item 14) vs Music element - Rhythm (Item 11) |
| No 5. | (iii) *‘Do the musical elements of waking audio counteract the effects of SI?’* | SI (Item 14) vs Music element - Tempo (Item 11) |
| No 6. | (iii) *‘Do the musical elements of waking audio counteract the effects of SI?’* | SI (Item 14) vs Music element - Pitch (Item 11) |
| No 7. | (iii) *‘Do the musical elements of waking audio counteract the effects of SI?’* | SI (Item 14) vs Music element - Volume (Item 11) |

Each test analysis was firstly trialled with a non-parametric contingency table analysis (Pearson’s Chi Square Test) to determine significant interactions between *SI* and each factor. Results of significance (*p* < 0.05) progressed for further analysis, while nonsignificant results (*p* > 0.05) where omitted from the study. Post-hoc testing included the generation of cell residuals (z*-*score) which were then calculated as *p*-values of cell significance. This allowed for the targeting of specific cells showing a number of frequencies which are significant against the report of *SI* intensity.

### Secondary analysis

For the significant (*p* < 0.05) results gathered from the primary analysis, we performed a secondary series of analyses. On this condition, we tested the appropriate items (10, 11, 13) against each significant result obtained. By conducting this sequence of analyses, we can respond to the main research questions by defining secondary level interactions between the primary results and each conditionally relevant item (I.e. sound type Item 10, subjective feeling Item 13, and music elements Item 11). These results provide data to be applied in the formulation and design of waking audio for *SI* in future studies. Each secondary analysis performed can be viewed in Table 4.

**Table 4.** Secondary Post-hoc analysis - Melody vs sound type and musical elements.

| TEST ANALYSIS | ANALYSIS COMPARISONS |
| --- | --- |
| No 8. | Music element - Melody (Item 11) vs Sound type (Item 10) |
| No 9. | Music element - Melody (Item 11) vs Music element - Rhythm (Item 11) |
| No 10. | Music element - Melody (Item 11) vs Music element - Tempo (Item 11) |
| No 11. | Music element - Melody (Item 11) vs Music element - Pitch (Item 11) |
| No 12. | Music element - Melody (Item 11) vs Music element - Volume (Item 11) |

Each secondary (Table 4) analysis was firstly tested with a non-parametric contingency table analysis (Pearson’s Chi Square Test) to determine significance against the conditional factors. Results of significance (*p* < 0.05) progressed for further analysis, while nonsignificant results (*p* > 0.05) where omitted. Further Post-hoc testing included the generation of cell residuals (z*-*score) which were then calculated as *p*-values of cell significance. This produced specific cells with significance against the conditional factors.

Each test analysis above (No. 8, 9) implemented the Fisher’s Exact Test utilising the Exact Monte Carlo method using 100000 repeated random samples, and the Post-hoc testing generated cell residuals (z_score) calculated as *p*-values of cell significance for each test analysis.

To conclude the analysis, the final three items of the questionnaire (Item 15, 16, 17, Fig 2) are tested against *SI* with a non-parametric contingency table analysis (Pearson’s Chi Square Test) to determine significance. These tests analyse whether a report for how long it takes to completely wake up (Item 15), the period after waking to complete the questionnaire (Item 16), and time of day completing the test (Item 17) influence the intensity of *SI*. See Table 6.

**Table 5.** Secondary analysis cases of significance.

| TEST ANALYSIS | ANALYSIS COMPARISONS |
| --- | --- |
| No 8. | Music element – Melody (Item 11) vs Sound type (Item 10) |
| No 9. | Music element - Melody (Item 11) vs Music element – Rhythm (Item 11) |

**Table 6.** Analysis No 13, 14 15.

| TEST ANALYSIS | ANALYSIS COMPARISONS |
| --- | --- |
| No 13. | $SI$ (Item 14) vs How long does it take you to 'come to' in the morning? (Item 15) |
| No 14. | $SI$ (Item 14) vs How long after waking did you complete this questionnaire? (Item 16) |
| No 15. | $SI$ (Item 14) vs At what time of day did you complete this questionnaire? (Item 17) |

## Results

Tables 7, 8, & 9 tabulate the counts and percentages of the raw data for the ‘Respondents’, ‘*Music appreciation & aptitude’*, and **‘***Alarm adoption & application’* demographic categories.

**Table 7.** Respondents.

| ITEM | CATAGORY | COUNT | % |
| --- | --- | --- | --- |
| Age group | 18 – 29 | 14 | 28 |
|  | 30 – 39 | 12 | 24 |
|  | 40 – 49 | 11 | 22 |
|  | 50 – 59 | 10 | 20 |
|  | 60+ | 3 | 6 |
| Gender | MALE | 15 | 30 |
|  | FEMALE | 35 | 70 |

**Table 8.** Music appreciation & aptitude.

| ITEM | CATAGORY | COUNT | % |
| --- | --- | --- | --- |
| Average music listening per day | 0-30 minutes | 19 | 38 |
|  | 31 minutes - 1 hours | 17 | 34 |
|  | 1 - 3 hours | 9 | 18 |
|  | 3+ hours | 5 | 10 |
| Music appreciation | Like a great deal | 31 | 62 |
|  | Like somewhat | 17 | 34 |
|  | Neither like nor dislike | 2 | 4 |
|  | Dislike somewhat | 0 | 0 |
|  | Dislike a great deal | 0 | 0 |
| Accuracy in describing music | Extremely well | 7 | 14 |
|  | Very well | 18 | 36 |
|  | Moderately well | 17 | 34 |
|  | Slightly well | 7 | 14 |
|  | Not well at all | 1 | 2 |

**Table 9.** Alarm adoption & application.

| ITEM | CATAGORY | COUNT | % |
| --- | --- | --- | --- |
| Device to play the audio | Clock Radio (alarm) | 7 | 14 |
|  | Mobile phone | 42 | 84 |
|  | Tablet | 1 | 2 |
|  | Laptop | 0 | 0 |
|  | Other | 0 | 0 |
| Days per week audio is used to wake up | 1. | 3 | 6 |
|  | 2. | 2 | 4 |
|  | 3. | 1 | 2 |
|  | 4. | 4 | 8 |
|  | 5. | 17 | 34 |
|  | 6. | 11 | 22 |
|  | 7. | 12 | 24 |
| Same audio to wake up every day | Yes | 46 | 92 |
|  | No | 4 | 8 |
| Volume increase over time | Yes | 17 | 34 |
|  | No | 33 | 66 |

### Primary analysis

Test analysis No 1: *SI* (Item 14) vs sound type (Item 10)

Test analysis No 2: *SI* (Item 14) vs subjective feeling (Item 13)

We found no significant interaction between the participants waking sound type and the reported intensity of *SI X*^*2*^ (12, *N* = 50) = 14.07, *p* = 0.296. Furthermore, the reported subjective feeling towards the waking sound and *SI* Fishers Exact Test returned no significant interaction *X*^*2*^ (*N* = 50) = 21.05, *p* = 0.066.

Test analysis No 3: *SI* (Item 14) vs musical element-melody (Item 11)

This test produced a significant interaction between the reported intensity of *SI* and the reported melodicity of the subjects’ waking sound *X*^*2*^ (16, *N* = 50) = 26.77, *p* = 0.044. The Fishers Exact Test result also produced a significant relationship between *SI* intensity and the melodicity of the waking sound with a value of *X*^*2*^ (*N* = 50) = 23.54, *p* = 0.022. The Post-hoc, Adjusted Residual testing of the data reports that there is a significant association between the intensity of *SI* as ‘Not at all’, and the reported waking sound melodicity as ‘Melodic’ (*Z* = 2.4, *p* = 0.022). Further, the intensity of *SI* is reported as ‘Often’ when compared to the waking sound melodicity as ‘Neither Unmelodic nor Melodic’ (*Z* = 2.4, *p* = 0.022).

### Secondary analysis

Test analysis No 8: Music element – melody (Item 11) vs sound type (Item 10)

Our results reveal a significant interaction between the reported waking sound melodicity, and the subjects’ reported waking sound type *X*^*2*^ (12, *N* = 50) = 24.38, *p* = 0.018. While Fisher’s Exact Test produced *X*^*2*^ (*N* = 50) = 28.99, *p* = 0.014. The Post-hoc, Adjusted Residual testing displayed several significant associations between the reported melodicity of the subjects waking sound in relation to the descriptions of the sound type. See Table 10.

**Table 10.** Secondary Post-hoc analysis for adjusted residuals of music element - melody vs sound type.

| TEST ANALYSIS No 8. RESULTS | Z score | p-value |
| --- | --- | --- |
| Unmelodic vs Alarm Tone | $Z = 2.9$ | $p = 0.005$ |
| Melodic vs Natural Sounds | $Z = 2.4$ | $p = 0.022$ |
| Very Melodic vs Musical Song | $Z = 2.1$ | $p = 0.044$ |

Test analysis No 9: Music element - melody (Item 11) Music element - rhythm (Item 11) There was a significant interaction between the reported melody of the waking sound, and the reported waking sound rhythm *X*^*2*^ (16, *N* = 50) = 50.32, *p* < 0.001. Subsequent analysis (Fisher’s Exact Test) produced *X*^*2*^ (*N* = 50) = 37.38, *p* < 0.001. The Post-hoc, Adjusted Residual testing revealed several significant interactions between the reported melody of the waking sound and rhythm of the waking sound. These results are shown in Table 11.

**Table 11.**
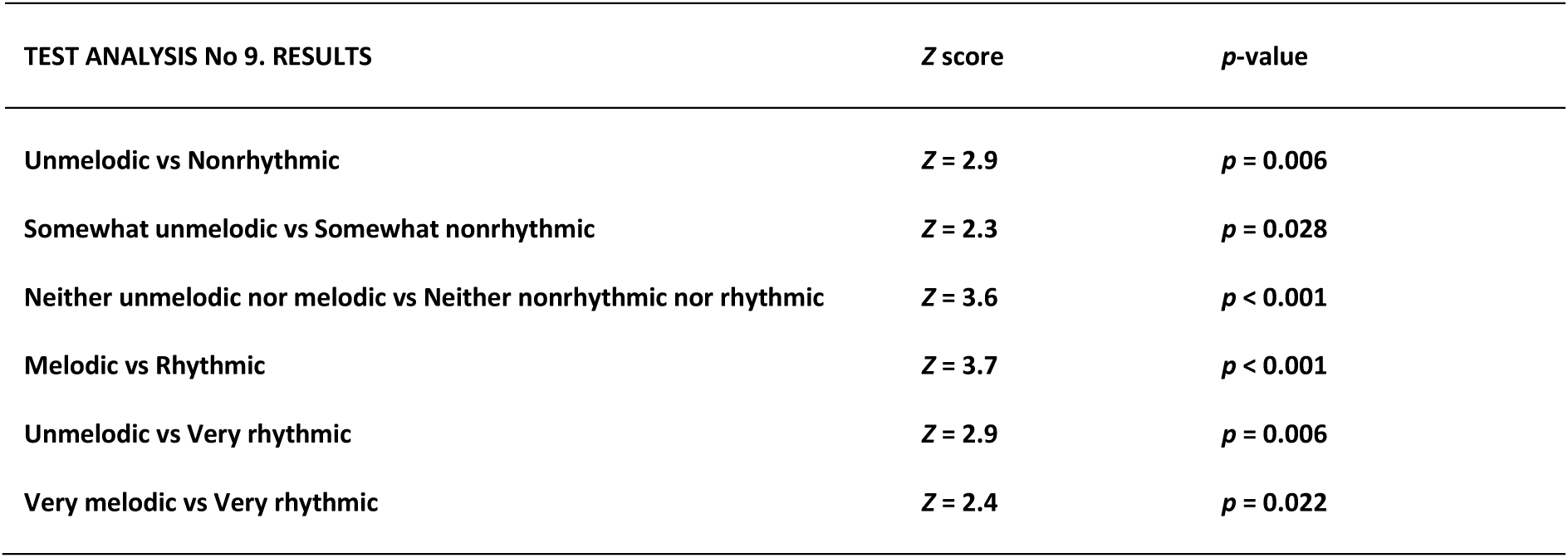
Secondary Post-hoc analysis for adjusted residuals of melody vs rhythm.

Test analysis No 13: *SI* (Item 14) vs How long does it take you to *‘come to’* in the morning?

Test analysis No 14: *SI* vs How long after waking did you complete this questionnaire?

Test analysis No 15: *SI* vs At what time of day did you complete this questionnaire?

No significant statistical interaction between these three cases and the intensity of *SI* were reported. See Table 12.

**Table 12.** Test analysis No. 13, 14, 15 results.

| TEST ANALYSIS | RESULTS |
| --- | --- |
| No 13. *SI* (Item 14) vs How long does it take you to '*come to*' in the morning? (Item 15) | $\chi^2 (N = 50) = 12.67, p = 0.124$ |
| No 14. *SI* (Item 14) vs How long after waking did you complete this questionnaire? (Item 16) | $\chi^2 (N = 50) = 25.23, p = 0.066$ |
| No 15. *SI* (Item 14) vs At what time of day did you complete this questionnaire? (Item 17) | $\chi^2 (N = 50) = 13.05, p = 0.110$ |

## Discussion

This research is the first experimental online questionnaire to analyse waking audio with respect to *SI*. Undertaken with a focus on reducing interventions to the participants and their usual waking routine, this research takes advantage of remote testing technologies to gather data in such a manner that was previously unattainable. The analysis present new insights to assist in the development and understanding of waking sound for *SI* reduction, and extends existing research in the field of auditory countermeasures for *SI*. Considering the increasing demands in this 24-hour society, maximising human alertness by reducing *SI* can assist in this evolution, providing a safer environment in situations where performance is critical.

Existing research supports audio’s potential to counteract the intensity of *SI* [11, 19]. In this research context, ‘excitative popular music’ has been shown to negate *SI* [19] which provides the first insight into a sound types effect on *SI*, though many questions concerning the specific auditory mechanics and aesthetics required for the design and production of such stimuli remain. The present study investigates this subject through a mixed methods self-report questionnaire, where an ecologically valid approach allows participants to submit responses from their preferred environment with minimal intervention. Through the questionnaire design and data obtained, we analyzed three primary research questions: (i) ‘Do waking sound types counteract the effects of SI?’, (ii) ‘Do subjective feelings towards waking sound types counteract the effects of SI?’, and (iii) ‘Do the musical elements of waking sound counteract the effects of *SI*?’. In response to these primary research questions we discuss the key findings, secondary analysis results, and response time evaluation and alertness factors.

From the analysis specific to the key research questions: (i) ‘Do waking sound types counteract the effects of *SI*?’, and (ii) ‘Do subjective feelings towards waking sound types counteract the effects of *SI*?’, our results indicate that reported waking sound types did not counteract *SI* intensity for our subject pool, nor did respondents’ subjective feelings towards their waking sound type. With respect to key research question: (iii) ‘Do the musical elements of waking audio counteract the effects of *SI*?’, the results show a significant interaction between the reported melodicity of waking audio and *SI*. Specifically, *SI* is evaluated to be neutralized (reported as ‘Not at all’) when analyzed against waking audio described as ‘melodic’ (See Results, Test analysis No 3: *SI* (Item 14) vs musical element-melody (Item 11)). Conversely, waking stimuli reported as ‘neither unmelodic nor melodic’ ‘often’ produce symptoms of SI (See results, Test analysis No 3: *SI* (Item 14) vs musical element-melody (Item 11)). These results indicate that waking audio perceived as ‘melodic’, irrespective of sound type, increase the probability of completely counteracting *SI*. By contrast, as the melodicity of waking stimuli decreases to a neutral interpretation (‘neither unmelodic nor melodic’), *SI* intensifies. Therefore, in response to primary research question (iii), ‘Do the musical elements of waking sound counteract the effects of *SI*?’, we conclude that musical elements may have the capacity to counteract *SI*, particularly waking audio interpreted as ‘melodic’, as this element has been analyzed to completely prevent *SI*.

To rationalize this phenomenon is challenging considering both the lack of specific research in this context, and the absence of descriptive detail (including music elements) concerning auditory stimuli for waking. Research shows that sound can increase and maintain arousal, and attract human attention [5–11], however, music elements (specifically melody) in the context of counteracting *SI* is unknown. We hypothesize that stimuli perceived as melodically neutral may be interpreted as an auditory ambient variation of their counterpart (melodic). When compared to strong melodic material this neutral classification is less likely to gain the human center of attention, may induce less arousal, and lead to reduced cognition, all of which, are symptoms of SI [2]. For situations where a stimuli’s melodic content is increased, its auditory ambience transitions to salience, increasing arousal, cognition and attention, subsequently reducing the effects of *SI*. For this analysis and building on available research, we propose that the melodic content of waking audio may be an essential musical factor in counteracting *SI*, supplementing holistic waking sound types as previously understood. Further, this result supports the requirement for detailed descriptions and inquiry of auditory test stimuli, to inform analysis and discussion in this research field.

Given the finding that melodic content of waking audio can influence *SI*, we now discuss this musical element and its statistical significance to waking sound types (e.g. alarm tone, natural sounds). Granting that *SI* is not directly affected in these conditions, we define which categories of waking sound types may have the ability to influence *SI* on a secondary level.

Our secondary analysis results show that an ‘unmelodic’ waking sound has a significant interaction to the category ‘alarm tone’, a ‘very melodic’ report is significant to the category ‘musical song’, and, ‘melodic’ is significant as a musical element ranking with respect to ‘natural sounds’ (See results, Test analysis No 8. Music element - Melody (Item 11) vs Sound type (Item 10)). From these results, we hypothesize that the musical mechanics and aesthetics of each sound type attributes to its perceived melodicity ranking. For example, as a function of the traditional sounding alarm (to wake sleeping humans through auditory intervention), by design this category of stimuli typically consists of a static rhythm, an insistent tonal center, and a salient aesthetic (e.g. a relentless beeping sound) [7, 8]. ‘Musical song’ incorporates melody as a dominant feature which is evident in popular western music, examples include, yet are not limited to, The Beach Boys ‘Good Vibrations’ [36], and The Cures ‘Close to me’ [37]. The relationship between ‘melodic’ and ‘natural sounds’ is more difficult to define. One hypothesis we present is bird song at dawn (a ubiquitous ecological waking sound common in many cultures [7]) for their melodic qualities associated to a variety of species [38], and by virtue that birds typically exist in natural environments. Similarly to the investigation of music elements and sound types in this study, bird song is categorized through song ‘type’ and the ‘elements’ which articulate the song type. [39].

The resulting interactions between melodicity and waking sound types may prove beneficial for the initial formulation of affective waking stimuli. We have shown that the melodicity of waking stimuli can counteract *SI*, and as melody is a musical element of significance with respect to particular waking sound types, we suggest that these factors must be carefully considered when designing optimal sounds to counteract *SI*. For instance, and referring to the results we have obtained, future investigation may study the ‘melodic’ element of ‘natural sound’ types for counteracting *SI*.

The quantity of rhythmic content in waking sound stimulus has a consistent and significant interaction to the perception of the reported melodicity (positive or negative) of waking audio. Our results show that when the rhythmic content of the waking stimuli is decreased, the reported melodicity is reduced (Table 11). In contrast, as the rhythmic content of the waking sound is increased, the melodicity of the stimuli is perceived in this manner also (Table 11). Additionally, the data reports an anomaly between the interpretation of ‘very rhythmic’ and melodicity. Specifically, if the rhythmic content of the waking sound is ‘very rhythmic’, then the perceived melodicity of the stimuli is reported as ‘unmelodic’ and ‘very melodic’ with equal probability (See Table 11). We hypothesize that this result is a by-product of the composition’s increased rhythmic elements, whereby a perceptual threshold is introduced, rendering the melodic and rhythmic interactions of the composition musically too complex for humans to clearly define as ‘very melodic’ or ‘unmelodic’. These results align with Boltz’s [40] findings suggesting an intrinsic relationship between melodic and rhythmic perception whereby temporal microstructures of compositions hold a primary role in the cognitive processing of melodic relations.

With respect to time duration evaluation of alertness, time period after waking to respond, and time of day with which the questionnaire was completed, there is no statistical interaction between these classes and *SI* (See Results, Test analysis No. 13, 14, 15). Our results indicate that *SI* intensity is independent of these factors highlighting the potential for waking stimuli to be as affective regardless of *SI* duration and time of day. Positive implications in this context include waking stimulus for shift workers, emergency response personal, and drowsy driving. However, further research is required in context to reinforce these understandings. For example, a Psychomotor Vigilance Test [41] could be utilized in conjunction with the SIQ in response to test stimuli to substantiate the results obtained and allow comparisons. Primarily a technical limitation in the context of online testing, this obstacle is being minimized with the increasing availability of accessible software to accomplish this task. Gorilla [23] is one online software system enabling the design and production of online questionnaires with embedded interactive capabilities and data logging.

The demographic data collected from the respondents in this study produced two key findings for referencing in waking sound development. Firstly, a mobile phone is the most frequently reported device for communicating the participant’s waking sound (84%, Table 9). Coupling this with estimates reporting that the audible frequency range of these devices ranges between 900 Hz – 10,000 Hz [42], this is central when developing auditory stimuli, as the human auditory hearing range spans 20Hz – 20,000 Hz [18]. Secondly, when considering the design of waking stimuli for *SI* and the aesthetic treatment of volume, sixty-six percent (66%) of subjects’ report that they employ a constant volume for waking. Specific research in this domain would clarify the benefits of auditory design targeting the most appropriate aesthetic volume treatment for waking sounds in the context of *SI*.

The primary analysis showed a significant statistical interaction between melodicity and *SI*. By dissecting the musical elements of the waking sounds melodic content in the secondary analysis, we conclude that the rhythmic attributes of the audio have a significant interaction to the perceived melodicity of the stimulus, which may impact *SI* through elemental musical interactions. We define these as key components to examine in the understanding of music and how its elements in combination can be assembled to produce the most effective waking sound stimuli to counteract *SI*. Future research based on these results may investigate rhythmic interactions with the perception of ‘melodic’ musical elements to inform the design and development of ‘natural sound’ types for *SI*.

## Supporting information

S1. Fig 1. Waking sound & sleep inertia questionnaire Section 1

S2. Fig 2. Waking sound & sleep inertia questionnaire Section 2

## Supporting information

S 1. Fig1. Waking sound & sleep inertia questionnaire Section 1.

S 2. Fig 2. Waking sound & sleep inertia questionnaire Section 2.

## Author contribution (s)

SJM and DSV designed the experiment; SJM collected the data; SJM, JEG and AGD analyzed the data; all authors edited and revised the manuscript and approved the manuscript for final submission.

## Author approvals

All authors have seen and approved the manuscript which has not been accepted or published elsewhere.

## Ethics statement

All research methods and data collection were approved by RMIT’s University College Human Ethics Advisory Network (CHEAN) (ref: CHEAN A 201710-02-17). Informed consent was given online by all participants by way of participating in the study.

## Data availability, DOI

10.6084/m9.figshare.7941128

