## Supplementary figures and images for "Alarm tones, music and their elements: A mixed methods analysis of reported waking sounds for the prevention of sleep inertia"

### S1. Fig 1. Waking sound & sleep inertia questionnaire Section 1

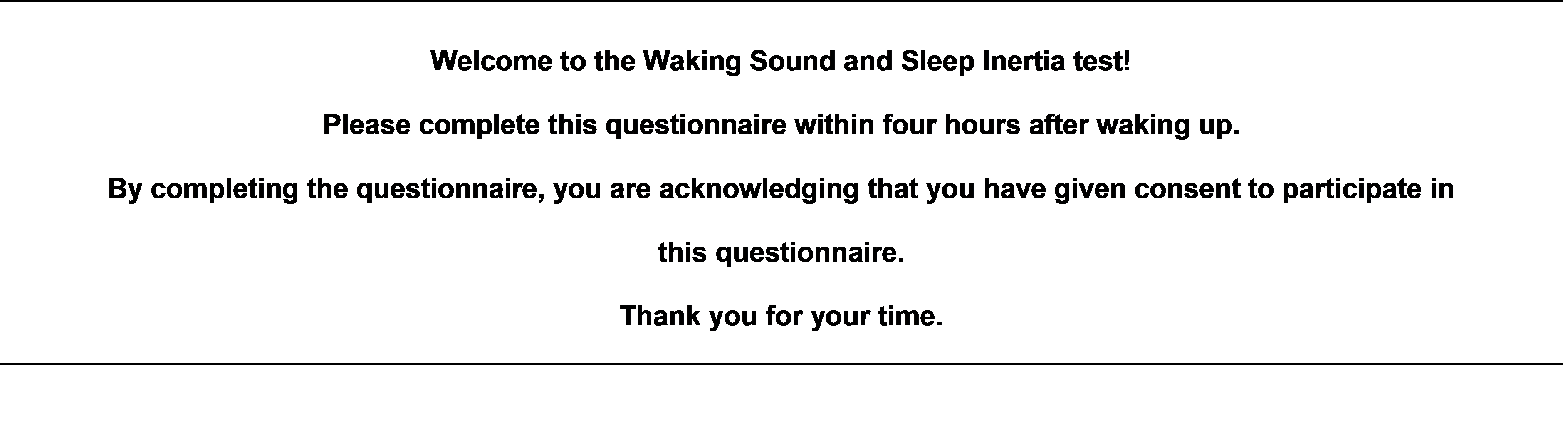

### S2. Fig 2. Waking sound & sleep inertia questionnaire Section 2

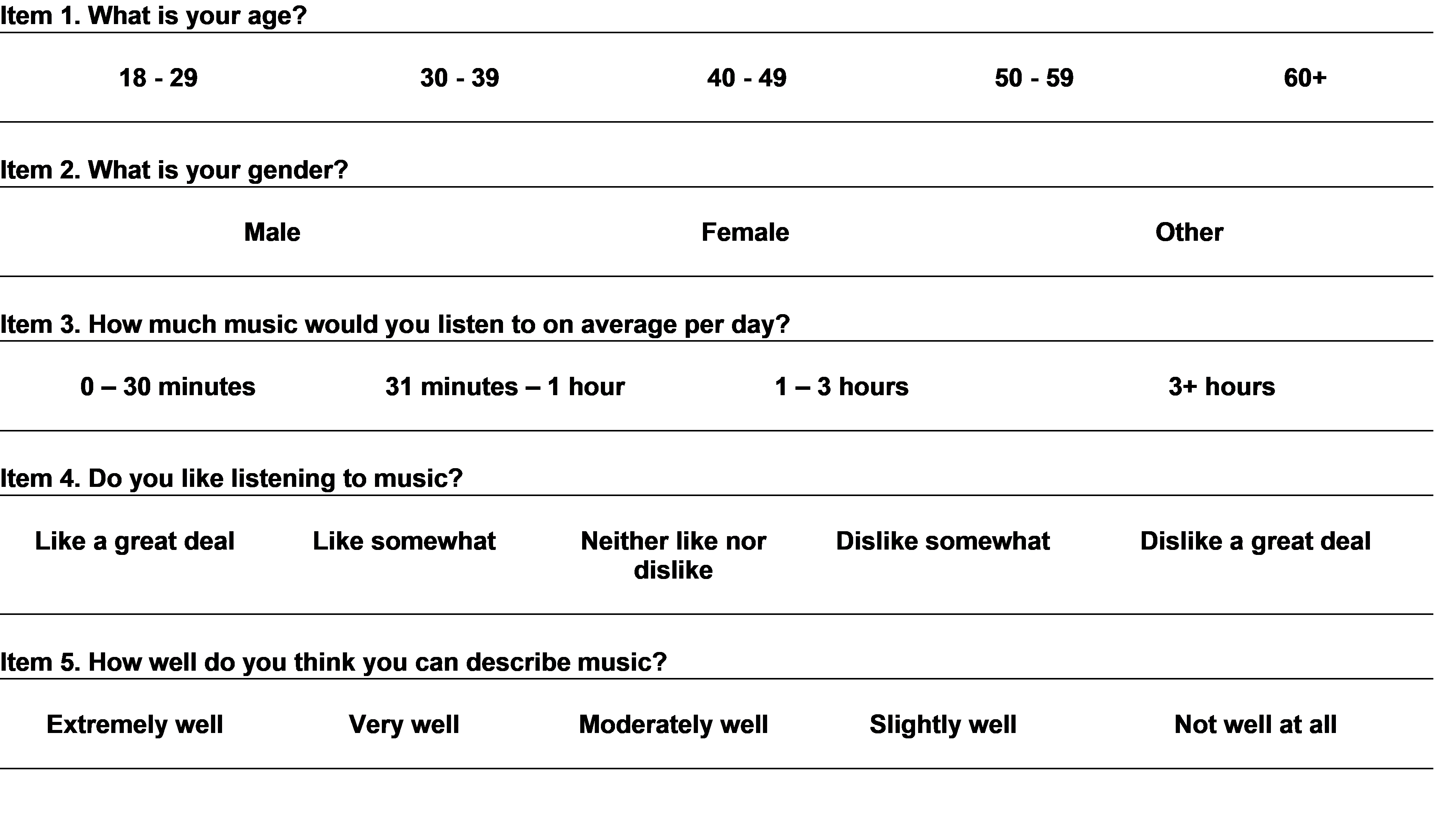
